# Hybrid capture-based sequencing enables unbiased recovery of SAR-CoV-2 genomes from fecal samples and characterization of the dynamics of intra-host variants

**DOI:** 10.1101/2020.07.30.230102

**Authors:** Yi Xu, Lu Kang, Zijie Shen, Xufang Li, Weili Wu, Wentai Ma, Chunxiao Fang, Fengxia Yang, Xuan Jiang, Sitang Gong, Li Zhang, Mingkun Li

## Abstract

**Background:** In response to the current COVID-19 pandemic, it is crucial to understand the origin, transmission, and evolution of SARS-CoV-2, which relies on close surveillance of genomic diversity in clinical samples. Although the mutation at the population level had been extensively investigated, how the mutations evolve at the individual level is largely unknown, partly due to the difficulty of obtaining unbiased genome coverage of SARS-CoV-2 directly from clinical samples.

**Methods:** Eighteen time series fecal samples were collected from nine COVID-19 patients during the convalescent phase. The nucleic acids of SARS-CoV-2 were enriched by the hybrid capture method with different rounds of hybridization.

**Results:** By examining the sequencing depth, genome coverage, and allele frequency change, we demonstrated the impeccable performance of the hybrid capture method in samples with Ct value < 34, as well as significant improvement comparing to direct metatranscriptomic sequencing in samples with lower viral loads. We identified 229 intra-host variants at 182 sites in 18 fecal samples. Among them, nineteen variants presented frequency changes > 0.3 within 1-5 days, reflecting highly dynamic intra-host viral populations. Meanwhile, we also found that the same mutation showed different frequency changes in different individuals, indicating a strong random drift. Moreover, the evolving of the viral genome demonstrated that the virus was still viable in the gastrointestinal tract during the convalescent period.

**Conclusions:** The hybrid capture method enables reliable analyses of inter- and intra-host variants of SARS-CoV-2 genome, which changed dramatically in the gastrointestinal tract; its clinical relevance warrants further investigation.

## Introduction

The ongoing Coronavirus disease 2019 (COVID-19) pandemic has brought a severe threat to public health and the global economy. The causative pathogen, severe acute respiratory syndrome coronavirus 2 (SARS-CoV-2) is likely of zoonotic origin. Genomes of RNA viruses mutate fast and undergo rapid evolution, which could ultimately affect their virulence, infectivity, and transmissibility. Monitoring variations in SARS-CoV-2 genome at the population level is vital for tracing the outbreak origin, tracking transmission chains, and understanding viral evolution (1–3). Meanwhile, the study of intra-host single nucleotide variation (iSNV), which represents the intermediate stage between the origin and the fixation of the mutation at the individual level, is essential for understanding how the virus evolves in the human body under the immune pressure (4). These studies require complete, in-depth, and unbiased profiling of SARS-CoV-2 genomes from a large number of patients.

However, there are still challenges in obtaining high-quality SARS-CoV-2 genome directly from clinical samples, especially for those with low viral loads. For samples with Ct > 30, the proportion of genome recovered was lower than 90% irrespective of the amplification and sequencing approach used (2). Current strategies for targeted enrichment of SARS-CoV-2 genome include hybrid capture and multiplex PCR amplification; the former method was proposed to be more reliable in generating unbiased coverage across the genome and identifying minor alleles (5). As the effectiveness of the hybrid capture method is influenced by multiple factors, such as probe design, rounds of hybridization, and viral load, the fidelity of hybrid capture warrants further evaluation in the context of iSNV investigation.

Recent studies have revealed the gastrointestinal (GI) tract as an important actor in the pathogenesis and transmission of COVID-19. GI infection of SARS-CoV-2 has been confirmed by the isolation of viable virus from the fecal specimen (6,7), and diarrhea is observed in > 20% of hospitalized COVID-19 patients (8). Moreover, the shedding of SARS-CoV-2 from the GI tract lasted much longer than that from the respiratory tract in children (9), raising the possibility of fecal-oral transmission. However, neither the genomic diversity of the virus nor its longitudinal dynamics in the GI is known.

To disentangle how SARS-CoV-2 evolves and adapts in the GI tract, 18 longitudinal fecal specimens from nine children with COVID-19 were collected. Nucleic acids of SARS-CoV-2 were enriched by a hybrid capture method. Seventeen near-complete genomes (> 99% genome covered) of the SARS-CoV-2 were obtained. Among them, the majority rule consensus sequences at four sites were changed in 1-5 days, and another 15 sites showed allele frequency changes greater than 0.3, indicating a rapid genomic change in the GI.

## Methods

### Sample collection and ethics

Eighteen fecal samples were collected from nine hospitalized children with real-time RT-PCR (RT-qPCR) confirmed SARS-CoV-2 infection. Two fecal samples were collected from each child within a time interval of 1-5 days (T1 and T2) in the recovery stage. The patients were characterized by mild symptoms and long duration of SARS-CoV-2 shedding in their feces (see Supplementary Table 1 for demographic and clinical information). All samples were inactivated at 56 □ for 30 min, and stored at −80 □ before processing.

This study was approved by the ethics review committee of Guangzhou Women and Children’s Medical Center. Written informed consents were obtained from the legal guardians of all children.

### Metatranscriptomic and hybrid capture-based sequencing

Each fecal sample (∼200 mg) was suspended in 1 mL PBS, and centrifuged at 8,000 g for 5 min to obtain the supernatant. RNA was extracted from 200 uL supernatant (AllPrep^®^ PowerViral^®^ DNA/RNA Kit, Qiagen), concentrated (RNA Clean & Concentrator™-5 with DNase I, Zymo Research), and used for library preparation (Trio RNA-Seq, Nugen). An aliquot of 750 ng library from each sample was used for hybrid capture-based enrichment of SARS-CoV-2 with one or two rounds of hybridization (1210 ssRNA probes, TargetSeq^®^ One nCov Kit, iGeneTech). Sequencing was performed on Illumina HiSeq X Ten platform. The load of SARS-CoV-2 was quantified with RT-qPCR targeting ORF1ab gene (Real-Time Fluorescent RT-PCR Kit for Detecting 2019-nCoV, BGI).

### Sequencing data analysis

Quality control of the sequencing reads including adapter trimming, low-quality reads removal, and short reads removal was done by fastp v0.20.0 (10) (-l 70, -x, -cut-tail, -cut_tail_mean_quality 20). Clean reads were mapped to SARS-CoV-2 reference genome Wuhan-Hu-1 (GenBank MN908947.3) using BWA mem v0.7.12 (11), followed by duplicate reads removal with Picard v2.18.22 (http://broadinstitute.github.io/picard). Mpileup files were generated with samtools v1.8 (12). Intra-host variants were identified using VarScan v2.3.9 and an in-house script (13). Criteria for variants included: (1) sequencing depth ≥ 50, (2) minor allele frequency ≥ 5%, (3) minor allele frequency ≥ 2% on each strand, (4) minor allele counts ≥ 10 on each strand, (5) strand bias of the minor allele < 10 fold, (6) minor allele was supported by the inner part of the read (excluding 10 base pairs on each end), and (7) minor allele was supported by ≥ 10 reads that were classified as *Betacoronavirus* by Kraken v2.0.8-beta (14) on each strand.

The ratio of the number of non-synonymous substitutions per non-synonymous site (Ka) to the number of synonymous substitutions per synonymous site (Ks) was calculated using KaKs_Calculator 2.0 (15) with the MS method.

### Data availability

Viral reads were deposited in the Genome Warehouse in the National Genomics Data Center (16) under project PRJCA2828, which are publicly accessible at https://bigd.big.ac.cn/gsa.

## Results

### Hybrid capture effectively enhanced SARS-CoV-2 reads and genome coverage

Eighteen fecal samples were collected from nine children with COVID-19, two samples were collected from each child within a time interval of 1-5 days (T1 and T2). Ages of the children ranged from three months to 13 years old. Their symptoms were either mild or moderate, and two of them had diarrhea. All samples were collected during the recovery stage (see Supplementary Table 1 for more details). We performed metatranscriptomic sequencing (Raw) and hybrid capture-based sequencing (with one-round of hybridization, E1) on all samples. Meanwhile, to investigate the efficiency of hybridization, nine samples were processed with an extra round of hybridization (E2). Two fecal samples from two healthy children were used as negative controls (NC). More than 25 million reads were generated for each sample except that no less than 12 million reads were generated for each NC sample. For COVID-19 samples, Ct values of RT-qPCR targeting SARS-CoV-2 varied from 28 to below detection limit, with a median of 33.26.

A median number of 157 (range 0-20,816) reads per million (RPM) reads were mapped to the SARS-CoV-2 reference genome in Raw data. As expected, the number of SARS-CoV-2 reads was negatively correlated with the Ct value (Spearman r = −0.59, p < 0.01, Fig 1A). One-round hybrid capture greatly improved the number of SARS-CoV-2 reads in all samples (RPM range 136-763,746, median 82,137, p < 0.0001, Wilcoxon matched-pairs signed rank test). An extra round of hybridization enabled further improvement (RPM 47,072-810,801, 718,025, p < 0.01), which was 19,726 and 33 times higher than those in Raw and E1 data, respectively (Fig 1A). Consequently, the hybrid capture method significantly increased SARS-CoV-2 genome coverage (Fig 1B-C, Table 1). In addition, we detected 24 and 12 SARS-CoV-2 RPM in two NC samples with one-round hybrid capture, indicating that the signal of SARS-CoV-2 observed in the COVID-19 patients was much stronger than that in NC.

**Figure 1.**
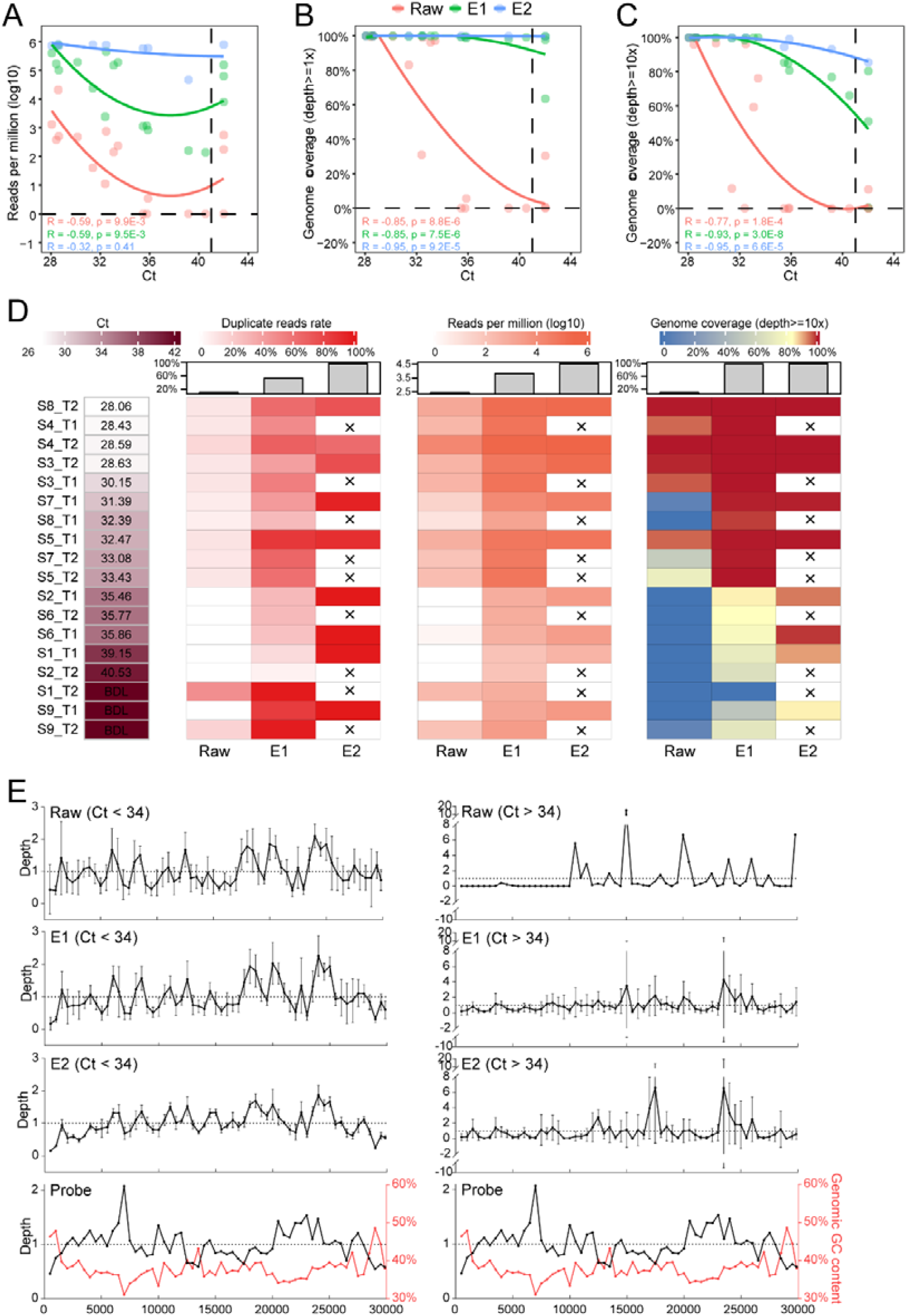
SARS-CoV-2 read counts and genome coverage obtained using direct metatranscriptomic sequencing and hybrid capture-based sequencing. (A) The number of SARS-CoV-2 reads in the unit of reads per million (RPM). (B-C) Genome coverage with depth ≥ 1 (B) and depth ≥ 10 (C). In A-C, local polynomial regression line, R and p values of Spearman correlations are shown; Ct value that below the detection limit was replaced with 42 for better visualization. (D) Heatmap of Ct, duplicate reads rate, RPM, and genome coverage after de-duplication. The median number for each group is shown with a bar plot, and crosses indicate that the samples were not included in E2 data. BDL, below the detection limit. (E) Depth distribution along SARS-CoV-2 genome. The relative depth was calculated for every 500 bases, and normalized to the average depth of each sample or the probes; only samples with average depth > 1 are shown. Red curve indicates GC content along the reference genome.

**Table 1.** Coverage of SARS-CoV-2 genome obtained with direct metatranscriptomic sequencing and hybrid capture-based sequencing.

| Group/Coverage<br>(range,<br>median, %) | Before de-duplication |  |  | After de-duplication |  |  |
| --- | --- | --- | --- | --- | --- | --- |
| | Depth $\geq 1$ | Depth $\geq 10$ | Depth $\geq 50$ | Depth $\geq 1$ | Depth $\geq 10$ | Depth $\geq 50$ |
| Raw | 0.0-100.0,<br>57.0 | 0.0-100.0,<br>11.4 | 0-99.9,<br>0.9 | 0.0-100.0,<br>57.0 | 0.0-100.0,<br>9.2 | 0-99.9,<br>0.4 |
| E1 | 63.5-100.0,<br>99.9 | 0.3-100.0,<br>99.1 | 0.2-100.0,<br>94.6 | 63.5-100.0,<br>99.9 | 0.3-100.0,<br>98.9 | 0.1-100.0,<br>92.4 |
| E2 | 99.7-100.0,<br>99.9 | 85.5-100.0,<br>99.9 | 22.8-100.0,<br>99.7 | 99.7-100.0,<br>99.9 | 85.1-100.0,<br>99.7 | 18.1-100.0,<br>99.7 |

An excess number of PCR cycles were implemented to ensure enough input for sequencing following the hybridization due to the loss of non-target nucleic acids. Accordingly, we noted that the proportion of duplicated reads increased from 10.9% in the Raw to 53.1% after the first hybridization and 95.5% after the second hybridization (Fig 1D). Despite the higher proportion of duplicate reads, the valid number of SARS-CoV-2 reads and genome coverage after de-duplication still significantly increased after hybrid capture (E1 *vs*. Raw: median fold change 246.2; E2 *vs*. E1: 2.7, Fig 1D, Table 1). Through our calculation, for samples with Ct < 29, direct metatranscriptomic sequencing (with 6G data) is sufficient to obtain a complete viral genome with a sequencing depth of at least 10-fold. One and two rounds of hybrid capture are needed for samples with Ct 29-34 and Ct 34-39 to cover the complete genome, respectively. By contrast, it is challenging to obtain adequate genome coverage for samples with Ct > 39 even when using two-round hybrid capture.

Besides the sequencing coverage, the sequencing depth distribution is another critical metric for downstream analysis; thus, we examined whether the depth distribution is well-maintained after hybrid capture. For samples with Ct < 34, similar depth distributions were observed between the Raw data and capture enriched data (Raw *vs*. E1, R^2^ = 0.88, Raw *vs*. E2, R^2^ = 0.60, E1 *vs*. E2, R^2^ = 0.76, p values < 0.0001, linear regression, Fig 1E, Fig S1A-C). Of note, the depth distribution was only slightly influenced by the probe density and genomic GC content, as the correlation was relative low between the depth and probe density (E1, R^2^ = 0.01, E2, R^2^ = 0.03, p values < 0.01, linear regression, Fig S1D-F) and between the depth and GC content (E1, R^2^ = 0.02, p <0.001, E2, R^2^ = 0.10, p < 0.0001, Fig S1G-I). The standard deviation of depths among samples decreased from Raw to E1 (median 0.40 and 0.29, p < 0.001, Wilcoxon matched-pairs signed rank test), and further to E2 (median 0.14, p < 0.0001), demonstrating a consistent recovery of the viral genome with hybrid capture. For samples with Ct > 34, hybrid capture resulted in unexpected depth peaks (17000-17500 and 23500-24000) that could not be explained by the probe density or the GC content.

### Hybrid capture enabled reliable inter- and intra-host variants profiling

By comparing the consensus sequences (major allele frequency > 0.7) of E1/E2 to those of Raw, we identified 41 inconsistent sites (with depth > 5) (Supplementary Table 2). However, all discrepancies involved in intra-host variants (the frequency of the major allele in at least one dataset < 0.7), and only four of them involved in major allele switches with frequency changes varied from 0.17 to 0.54 (three sites with depth < 10), suggesting high reliability of genome obtained with hybrid capture.

We further compared the frequencies of alternative alleles (non-reference allele, AAFs) among Raw, E1, and E2. Although the number of mutations detected in Raw was much less than those in E1 (201 *vs*. 460) which may be caused by the limited genome coverage in Raw, there were decent linear correlations among AAFs of Raw, E1, and E2 (Raw *vs*. E1, R^2^ = 0.90; Raw *vs*. E2, R^2^ = 0.84; E1 *vs*. E2, R^2^ = 0.69; p values < 0.001, Fig 2A-C). Moreover, the changes of AAFs from T1 to T2 was also significantly correlated between the Raw and E1 (R^2^ = 0.20; p < 0.001, Fig 2D). Notably, the frequency change calculated from the Raw data showed a larger variance than E1 (Fig 2D), likely reflecting more significant stochastic fluctuations related to the lower sequencing depth. These results demonstrated that hybrid capture-based sequencing is ideal for genomic diversity analysis, and also enable an accurate longitudinal analysis of iSNVs.

**Figure 2.**
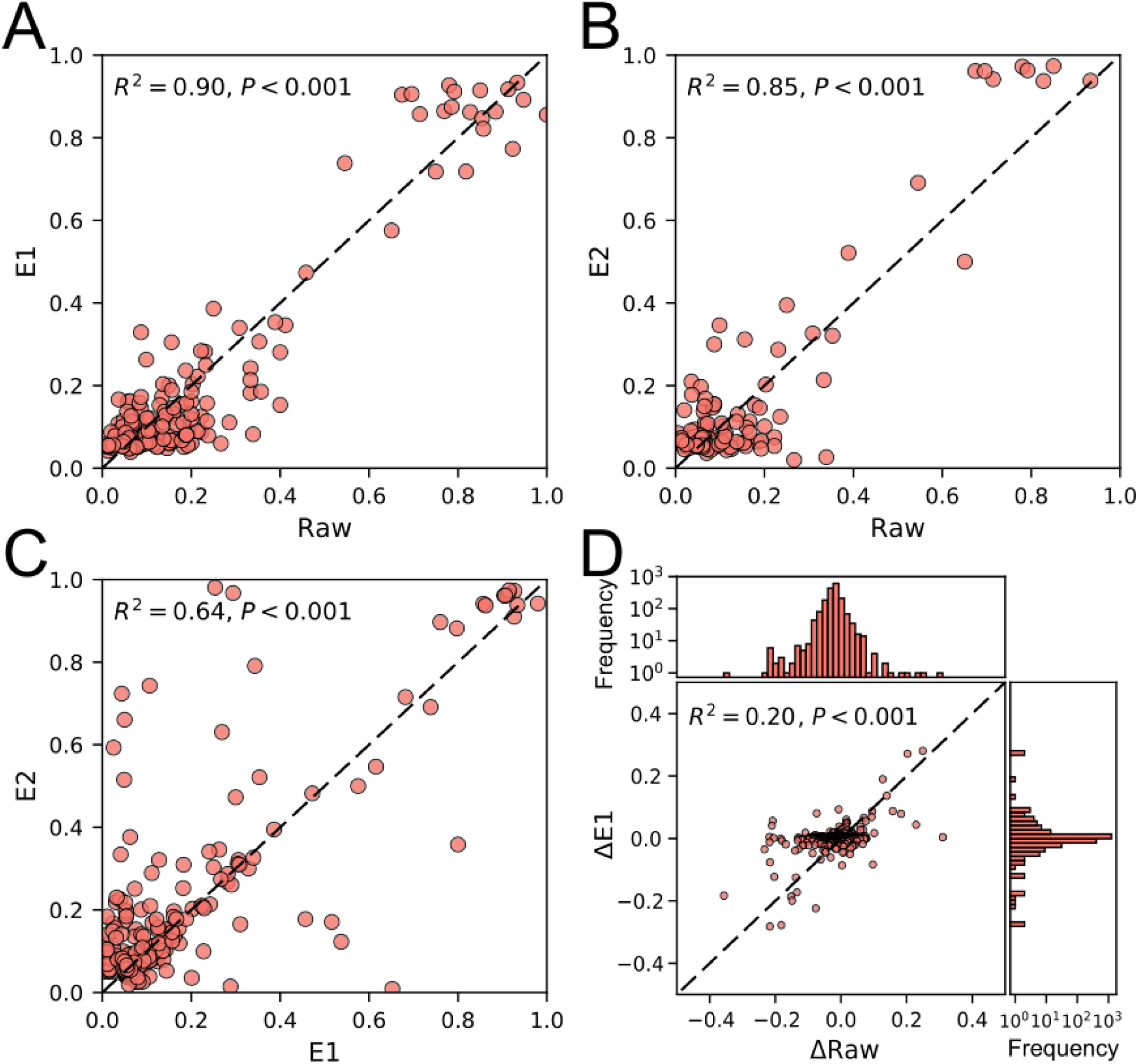
Evaluation of hybrid capture in mutant allele frequency profiling. (A-C) Consistency among alternative allele frequencies (AAF) in Raw, E1, and E2 data. (D) Consistency between AAF changes from T1 to T2 in Raw and E1 data. R^2^ and p values of linear regressions are shown.

### iSNV characteristics and longitudinal dynamics in fecal samples

Taking advantage of the higher sequencing depth and high fidelity of SARS-CoV-2 genome obtained using hybrid capture method, E1 and E2 data were pooled together for iSNV analysis. A total number of 229 iSNVs were identified at 182 sites (criteria were described in Methods), including 159 non-synonymous, 60 synonymous, 3 non-coding, and 7 stop-gain mutations. The overall Ka/Ks ratio was 0.54, which was significantly different from 1 (Fisher’s exact test, p < 0.001) and thus suggested a purifying selection. We noted that only 34% (78 out of 229) of the iSNV were polymorphic in the population (by comparing with 12063 mutations retrieved from https://bigd.big.ac.cn/ncov, 2020 Jun 9), suggesting that a large proportion of mutations cannot be fixed in the individual or transmitted to other individuals. However, the absence of the mutation in the population dataset could also be attributed to the limited genomic diversity included in the current population dataset. The number of mutations was inversely correlated with mutant allele frequency (MAF), except for iSNVs with MAF 0.8-0.95 (Fig 3A), which probably reflected the back mutation from the mutant allele to the ancestral allele. The number of iSNVs in different genes is proportional to the length of corresponding regions (Fig 3A), suggesting no visible mutation hotspot on the SARS-CoV-2 genome.

**Figure 3.**
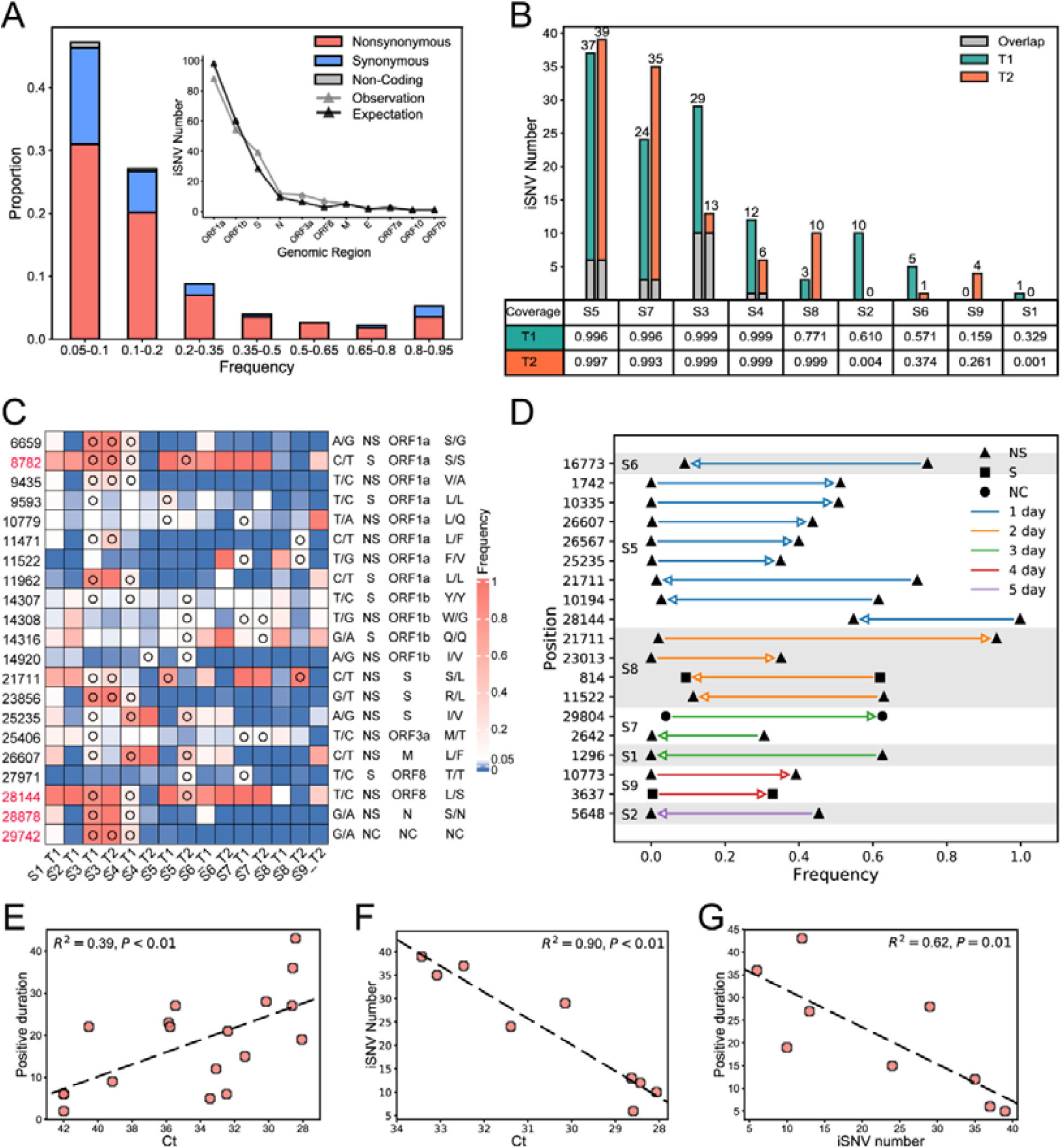
iSNV profiles and longitudinal dynamics in fecal samples. (A) Alternative allele frequency distribution of all iSNVs. The inserted plot shows the number of observed iSNVs in different genes; the expected number of iSNVs was calculated based on the length of each region. (B) The number of iSNVs identified in each sample. Genomic coverages with depth > 50 are shown below the plot. (C) AAF at iSNV positions shared by multiple individuals. The nucleotide at the position (reference allele/alternative allele), mutation type, genomic region, and amino acid change are shown on the right of the heatmap. Open circles indicate identified iSNVs. The four positions associated with recurrent mutations (3) are highlighted in red. (D) iSNVs with frequency change > 0.30 from T1 to T2. Only changes with adjusted p values < 0.05 (Fisher’s exact test) are shown. Arrows indicate the direction of changes from T1 to T2, and the colors of lines indicate time intervals between T1 and T2. (E-G) Correlation between Ct value, iSNV number, and the duration of detection of SARS-CoV-2 RNA in feces since T1/T2. R^2^ and p values of linear regressions are shown. In F and G, only samples having full genomic coverage with depth ≥ 50 were included (n = 9). NS, non-synonymous, S, synonymous, NC, non-coding.

The number of iSNVs markedly varied among different individuals, ranging from 6 to 39 in samples with full genome covered by at least 50-fold (Fig 3B). Twenty-one iSNVs were shared among multiple individuals, which is more than expected by chance (P<0.001, Exact Poisson test, Fig 3C). Moreover, 4 of the shared iSNVs were located at mutation hotspots (3). Considering that only 198 such mutation hotspots were identified, shared iSNVs were significantly enriched at positions with higher mutation rates (P<0.001, Fisher exact test). C8782T and T28144C are widespread variants that defined a subgroup of SARS-CoV-2 (17,18); they were both heterogeneous in multiple samples in our data, suggesting that either the position is prone to mutate or multiple strains were circulating in the population when the samples were collected. Mutations at most positions only involved transitions between two nucleotides, except that more than two nucleotides were observed at four positions, namely 14307, 14308, 11147, and 11148 (Fig 3C, Supplementary Table 3).

The dynamics of all iSNVs were then examined, and only 8.3%-34.5% of iSNVs were observed at both T1 and T2 (Fig 3B). Meanwhile, 19 iSNVs showed frequency changes greater than 0.30 within 1-5 days (Fig 3D), 16 of which are non-synonymous mutations. Nine of these significant shifts occurred within a day and four within two days, implying intense selection pressure or strong genetic drift. Notably, the most dramatically shifted iSNV C21771T, which caused a non-synonymous mutation from Ser to Leu, was observed in two individuals but showed opposite tendencies, whereby the frequency of the mutant allele increased by 0.91 in one day in S8 while decreased by 0.71 in two days in S5. Although we favored a random genetic drift hypothesis, considering that the viral genome differed at more than 20 positions between S8 and S5, the possibility of distinct selection pressures under different genomic backgrounds cannot be ruled out. Besides, the number of rapidly-changing positions varied among different individuals, which may reflect different immune pressures (Fig 3D). We also noted that some mutations in the same individual had similar frequencies and showed concurrent changes (S5 and S8), suggesting they were located on the same haplotype.

We further investigated the correlation between virological features and clinical features. First, the load of SARS-CoV-2 was found to be positively correlated with the duration of detection of SARS-CoV-2 RNA in feces since T1/T2 (linear regression, R^2^ = 0.39, p < 0.01, Fig 3E). Second, the number of iSNV was in a negative correlation with SARS-CoV-2 load (R^2^ = 0.90, p < 0.01, Fig 3F), as well as the duration of detection of SARS-CoV-2 RNA (R^2^ = 0.62, p = 0.01, Fig 3G). The increased genomic diversity in samples with lower viral loads may reflect a stronger immune response to the virus, which enables faster elimination of the virus in the body and thus associated with better clinical outcomes.

## Discussion

The limited viral load in clinical samples has long been an obstacle to both pathogen detection and genomic studies, especially considering that the abundance of viruses could decrease when the disease progresses. Here we have demonstrated that the hybrid capture method was capable of recovering unbiased SARS-CoV-2 genome from RT-qPCR-negative and metatranscriptomic sequencing-negative samples. Furthermore, it also enables reliable analysis of inter- and intra-host variants, making it a promising strategy for genomic studies on the SARS-CoV-2.

The identification of iSNVs in fecal samples as well as in throat swab and bronchoalveolar lavage fluid, suggest ongoing evolution of SARS-CoV-2 in the human body (19–22). However, the possibility of infection of multiple strains cannot be entirely ruled out. With the time series samples, we have proved that novel mutations could occur within one day, and the frequency of the mutation changed notably fast in humans. Thus, iSNVs are more likely to represent spontaneous mutations rather than infection of multiple strains. A recent study proposed that SARS-CoV-2 in GI had a relatively higher genomic diversity compared to that in the respiratory tract (20), suggesting that the mutation rate may vary among different tissues. Thus, the fast-evolving of the viral genome in GI observed in our study may not be applicable to other organs.

All samples in this study were collected from pediatric patients in rehabilitation with no clinical symptoms. There had been more than two weeks since symptoms onset for most patients except Sample 5. Fecal shedding of SARS-CoV-2 had been observed in multiple studies and was proposed to be longer compared to respiratory samples(9,23). However, whether the RNA shedding from stools is infectious during the convalescent phase is unclear(24). The evolving of the viral genome in feces observed in our study suggested that the virus was viable and actively replicated in the gastrointestinal tract. Thus we speculate that the fecal transmission of SARS-CoV-2 is possible, and further study is warranted to investigate whether the viral load in feces is sufficient to establish an infection.

Although a general purifying selection on the viral genome had been shown in our data and also by previous studies, we did not observe any signature of selection on specific positions. Wang et al. found increased frequencies of two mutations (C21711T and G11083T) in two samples, and suspected an adaptive selection on these mutations (20). Interestingly, C21711T was also observed in our data and showed the greatest frequency changes. However, instead of simultaneous increases, the direction of the frequency change was opposite in two samples, indicating a robust random drift effect. Of note, the viral genome of the two samples showed significant divergence (> 20 substitutions), and how the genomic background interacts with a single mutation needs further investigation.

C8782T and T28144C had been reported to be in linkage disequilibrium and used to define an ancestral subgroup of SARS-CoV-2 (18). The authors interpreted the heteroplasmy status at these two sites as evidence for multiple strains infection. However, our data indicated that these two positions were prone to mutate, and their frequencies were subject to change. Van Dorp et al. also identified these two positions as hotspots for recurrent mutations (3). Thus, the grouping of SARS-CoV-2 genomes based on these two mutations should be with great caution.

The mutation and their frequency change may also affect the response to drugs. Position 10194 locates in the region encoding 3-chymotrypsin-like cysteine protease (3CL^pro^), which is essential for the replication of coronavirus, and this region is also predicted to be the target for a few drugs including Lopinavir (25,26). At the first time point in patient S5, there were an equivalent number of reads presenting A and T at this position (a mutation from A to T results in an amino acid change from Glu to Val). At the second time point, which was one day later, the mutant allele T almost disappeared. Although it is unclear whether the mutation could affect the binding of anti-viral drugs, the risk should not be overlooked.

In summary, our study highlighted the need for extensive studies on the intra-host variant dynamics in different tissues, using large cohorts spanning a wide range of ages, disease severity, and geographic regions.

## Acknowledgments

We thank Dr. Xue Yongbiao and colleagues from National Genomics Data Center for helpful discussion and computational resource support. We thank Dr. Tang Hong and Dr. Liu Dongping from Institut Pasteur of Shanghai, Chinese Academy of Sciences for project coordination.

---

**Supplementary Figure 1. Correlation between the sequencing depth, depth distribution of probe, and genomic GC content**. (A-C) Correlation between depth of Raw, E1, and E2. (D-F) Correlation between depth of probe and depth of Raw, E1, E2, respectively. (G-I) Correlation between genomic GC content and depth of Raw, E1, E2, respectively. The relative depth was calculated for every 500 bases, and normalized to the average depth of each sample or the probes. Samples with Ct < 34 and average depth > 1 are shown.

